# Expression and purification of the mitochondrial transmembrane protein FAM210A in *Escherichia coli*

**DOI:** 10.1101/2023.05.27.542570

**Authors:** Jared Hollinger, Jiangbin Wu, Kamel M. Awayda, Mitchell R. O’Connell, Peng Yao

## Abstract

The protein Family with sequence similarity 210 member A (FAM210A) is a mitochondrial inner membrane protein that regulates the protein synthesis of mitochondrial DNA encoded genes. However, how it functions in this process is not well understood. Developing and optimizing a protein purification strategy will facilitate biochemical and structural studies of FAM210A. Here, we developed a method to purify human FAM210A with deleted mitochondrial targeting signal sequence using the MBP-His_10_ fusion in *Escherichia coli*. The recombinant FAM210A protein was inserted into the *E. coli* cell membrane and purified from isolated bacterial cell membranes, followed by a two-step process using Ni-NTA resin-based immobilized-metal affinity chromatography (IMAC) and ion exchange purification. A pulldown assay validated the functionality of purified FAM210A protein interacting with human mitochondrial elongation factor EF-Tu in HEK293T cell lysates. Taken together, this study developed a method for purification of the mitochondrial transmembrane protein FAM210A partially complexed with *E.coli* derived EF-Tu and provides an opportunity for future potential biochemical and structural studies of recombinant FAM210A protein.

## 1. Introduction

The mitochondrion is an essential organelle generating energy for most biological processes within eukaryotic cells. Within mitochondria, the electron transport chain (ETC) complex, located in the inner mitochondrial membrane (IMM), is responsible for the production of energy by creating an electrochemical-proton gradient across IMM that drives the synthesis of ATP [1]. Besides their function in energy production, mitochondria serve as functional hubs for many other important processes, such as metabolism, cell signaling, apoptosis, and calcium signaling [2]. These critical functions require the coordination of various proteins encoded by the nuclear genome or mitochondrial DNA. Mass spectrometry analysis of mitochondrial proteome from different tissues has identified >1000 different proteins in mammalian mitochondria [3], many of which are functionally unannotated. One major hurdle in studying these proteins in vitro is the difficulty in purifying them. Because of this hurdle, it is challenging to evaluate their biochemical properties, especially for IMM-localized mitochondrial transmembrane proteins.

The family with sequence similarity 210 member A (FAM210A) is a mitochondrial protein localized at the IMM. Human FAM210A protein contains 272 amino acids with multiple domains, including a mitochondrial targeting signal with a cleavage site at Val^95^, a domain of unknown function 1279 (DUF1279) containing a single transmembrane motif, and a coiled-coil domain at its C-terminus [4]. FAM210A is critical for embryonic development and viability in mice [5], and genome-wide association studies (GWAS) have found that mutations of FAM210A are associated with sarcopenia and osteoporosis in humans [6,7]. However, FAM210A is not expressed in bones but is most highly expressed in mice’s skeletal muscle, cardiac muscle, and brain, suggesting an important role in these mitochondria-enriched organs [7]. Although our previous study has shown that FAM210A is a potential mitochondrial protein translation regulator through binding to EF-Tu, a critical mitochondrial translation elongation factor [8], the biochemical properties of FAM210A protein and molecular mechanism in translational control remain elusive.

In this study, we have developed a protocol for the overexpression and purification of FAM210A in *Escherichia coli (E. coli)*. Human FAM210A, with a mitochondrial targeting sequence deletion (dMTS), was fused to an N-terminal MBP-His_10_ tag and overexpressed in BL21(DE3) *E. coli*. The fusion protein was successfully purified from isolated cell membranes, and the MBP-His_10_ tag was removed by a site-specific TEV protease allowing for the purification of the FAM210A-dMTS protein. Due to its high isoelectric point (pI), the purified FAM210A dMTS protein was further purified using ion exchange chromatography. As a functional assay, purified FAM210A protein was shown to pulldown human mitochondrial elongation factor EF-Tu from HEK293T cell lysates. These findings suggest that MBP-His_10_-FAM210A-dMTS protein can be successfully overexpressed in *E. coli*, where it is inserted into the cell membrane and then purified to allow for further biochemical assays.

## 2. Materials and Methods

### 2.1 PCR and molecular cloning

Two vectors with C-terminal MBP (maltose-binding protein)-His_6_ Tag (2Cc-T; Addgene Plasmid #55209) or N-terminal His_10_-MBP Tag (2CT-10; Addgene Plasmid #37237) were chosen as the MBP tag has been shown to increase the solubility of select proteins [9]. The full length (FL) or deletion of the mitochondrial targeting sequence (dMTS) of the human FAM210A coding sequence (CDS; Uniprot Entry # Q96ND0) region was amplified using Q5 DNA polymerase (NEB, Cat # M0491). The FAM210A DNA was inserted into 2Cc-T and 2CT-10 vectors using the Ligation Independent Cloning (LIC) method. The plasmids were linearized using the restriction enzyme *HpaI* for 2Cc-T and *SspI* for 2CT-10 at 37°C for 3 hours. The products of the PCR amplification for the insert and the digestion of the vectors were run on a 1% agarose gel and subsequently gel purified. The gel-purified plasmid and DNA insert were digested with T4 DNA polymerase and either dCTP or dGTP. If the insert DNA was digested in the presence of dCTP, the vector was digested in the presence of dGTP and vice versa. This reaction was conducted at room temperature for 30 minutes before heat inactivating the enzyme at 75°C for 20 minutes. The insert and vector with the appropriate overhangs were then placed together at room temperature for 30 min to allow for annealing. The plasmids were then transformed into DH5-alpha high-efficiency competent cells (NEB). Following transformation, correct cloning of the inserted sequence of FAM210A was confirmed via restriction digest and Sanger sequencing (Genewiz/Azenta).

The primer sequences for FAM210A FL and dMTS cloning are listed below:

210A-FL-2CCT-F: 5’ TTTAAGAAGGAGATATAGTTC ATGCAATGGAATGTACCACGGAC 3’

210A-FL-2CT10-F: 5’ TACTTCCAATCCAATGCA ATGCAATGGAATGTACCACGGAC 3’

210A-dMTS-2CCT-F: 5’ TTTAAGAAGGAGATATAGTTCATGTCATCCAGTGCCACAGCTCAG

GGA 3’

210A-dMTS-2CT10-F: 5’ TACTTCCAATCCAATGCA TCATCCAGTGCCACAGCTCAGGGA 3’

210A-2CCT-R: 5’ GGATTGGAAGTAGAGGTTCTC TTCCACTTTTTTCTTAAAGGAAAC 3’

210A-2CT10-R: 5’ TTATCCACTTCCAATGTTATTA TTCCACTTTTTTCTTAAAGGAAAC 3’

### 2.2 Overexpression of MBP-His_10_ tagged FAM210A fusion protein in *E. coli*

The expression vectors encoding for FL or dMTS FAM210A and MBP-His_10_ Tag fusion proteins were transformed into three different *E. coli* strains, BL21(DE3) (Thermofisher, Cat. # EC0114), Rosetta2 (DE3) (Novagen, Cat. #71397), and Lemo21 (DE3) (NEB, Cat. # C2528J) cells for initial overexpression trials. The cells were cultured overnight in 5 mL LB Broth with 5 μg/mL ampicillin. The overnight culture was used to inoculate a new 5 mL subculture. The subculture was grown until the optical density (OD) was 0.8. The cells were induced with various IPTG concentrations (GoldBio, Cat. # 2481C) at 37°C for 3.5 hours. 1 mL of induced culture was pelleted and resuspended in 200 μL of Protein Loading Buffer (National Diagnostics, Cat. # EC-887) before being heated at 95°C for 10 min. Each sample (10 μL) was loaded and separated on an SDS-PAGE followed by Coomassie G-250 staining (VWR, Cat. # 0615) to visualize the overexpression of the desired fusion protein.

### 2.3 Isolation and solubilization of bacterial cell membranes

Initial trials were carried out in 100 mL of culture with 100 μg/mL ampicillin. Induced cells were pelleted at 5,500 g for 12 min twice. The cell pellet was resuspended in Cell Lysis Buffer containing 50 mM Tris-HCl pH7.5, 250 mM NaCl, Protease inhibitor cocktail (Roche, Cat. # 11697498001), 1 mM PMSF, 1 mM DTT, and 5% glycerol. Resuspended cells were sonicated on ice for 10 sec, followed by 10 sec to cool down for 15 min. The suspension was cleared by centrifugation at 13,000 g for 15 min. The supernatant was transferred to ultracentrifuge tubes and spun at 160,000 g for 60 min to pellet the membrane fraction. The membrane pellet was resuspended in ice-cold PBS before being spun at 100,000 g for 60 min for the wash step. The pellet was resuspended in a solubilization buffer containing 2% n-Dodecyl-B-D-Maltoside (DDM, GoldBio, Cat. # DDM10), 350 mM NaCl in PBS. The resuspension was rotated gently at 4°C for 2.5 hours to solubilize the lipid bilayer. The solubilized mixture was then spun at 100,000 g for 1 hour to pellet any debris leftover leaving solubilized fusion protein in the supernatant. The supernatant was placed with equilibrated Ni-NTA (GoldBio, Cat. # H-350) resin and allowed to bind overnight.

### 2.4 Affinity purification by nickel-based immobilized metal affinity chromatography

The solution of bound resin was transferred to a gravity flow purification column (Bio-Rad, Cat. # 7311550). The flowthrough was collected, and the Ni-NTA resin was washed with 3 column volumes of wash buffer containing 50 mM Tris-Cl (pH 7.5), 500 mM NaCl, 10 mM imidazole, 5% glycerol, 1 mM DTT, and 0.2% DDM. The MBP-His_10_-FAM210A-dMTS fusion protein was eluted off the resin with an elution buffer. The elution buffer contained 500 mM NaCl, 50 mM Tris-Cl pH 7.5, 5% glycerol, 1 mM DTT, 0.2% DDM to maintain micelles, and 400 mM imidazole. The elution fraction was concentrated using an Amicon centrifugal filter with 10 kDa MWCO (molecular weight cut-off) (Millipore, Cat. # UFC8010). A bicinchoninic acid (BCA) assay was performed to calculate the overall protein concentration. Samples were collected at each step for analysis by SDS-PAGE gel to determine if the fusion protein was inside the membrane fraction.

### 2.5 Cleavage of MBP-His_10_ Tag by TEV protease

Prior to the addition of TEV protease, a buffer exchange was carried out using a 10 kDa MWCO centrifugal filter (Millipore, Cat. # UFC8010) to remove the imidazole present in the elution buffer. The cleavage reaction using TEV protease (Genscript, Cat. # Z03030) was subsequently carried out at 4°C for 16 hours. Equilibrated Ni-NTA resin was added to the products to bind the MBP-His tag and the His-tagged TEV protease (Genscript, Cat. # Z03030) while allowing the purified FAM210A-dMTS cleaved product to be collected in the flowthrough. TEV was added based on the quantity of the protein concentration that needed to be cleaved. We used 1 unit of the TEV protease per 3 μg of the fusion protein.

### 2.6 Ion exchange chromatography

A HiTrap HP SP column (Cytiva, Cat. # 17115201) and an AKTA pure FPLC system (Cytiva) were used to perform cation exchange chromatography to further purify FAM210A-dMTS. The column was initially equilibrated in 50 mM Tris-Cl pH 7.5, 5% glycerol, 1 mM DTT, 0.01% DDM, and 50 mM NaCl. FAM210A-dMTS was buffer exchanged into the same buffer immediately before the chromatography run using a 10 kDa MWCO centrifugal filter. A 50 mM-1M NaCl gradient in the same buffer was used to elute FAM210A-dMTS. Fractions were collected for further concentration and analysis.

### 2.7 Size exclusion chromatography (SEC)

Cleaved FAM210A-dMTS protein at a concentration greater than 1 mg/mL was injected onto a Superdex 200 Increase 10/300 GL gel filtration column (Cytiva, Cat. # 28990944) running on an AKTA pure FPLC system preequilibrated with size exclusion buffer (50 mM Tris-Cl pH 7.5, 250 mM NaCl, 1 mM DTT, 5% glycerol, 0.1% DDM). Fractions were analyzed with SDS-PAGE to examine the presence of FAM210A.

### 2.8 SDS-PAGE and Western blots

SDS-PAGE gels were prepared using Protogel 30% (National Diagnostics, Cat. # EC-890) at varying percentages of acrylamide ranging from 10-12%. All protein samples were prepared using 5x Loading Buffer (National Diagnostics, Cat. # EC-887). For Western Blots, the samples were separated on 10% freshly cast SDS-PAGE gels followed by a wet-transfer method using a 0.45 μm Immobilon-P PVDF membrane (Millipore, Cat. # IPVH00010). The transfer was performed in the prechilled 1x Tris-glycine transfer buffer (25 mM Tris, 190 mM glycine, 20% v/v methanol) for 1-1.5 hours. The membrane was then placed into a blocking buffer containing 5% milk in Tris-buffered saline with 0.1% Tween-20 (TBS-T). All primary antibodies were prepared in TBS-T containing 4% (w/v) bovine serum albumin (BSA). Rabbit polyclonal anti-FAM210A antibody (HPA014324; IB: 1:1000) was purchased from Sigma-Aldrich. Mouse monoclonal anti-EF-Tu antibody (sc-393924; 1:500) was purchased from Santa Cruz. The secondary antibodies used in the present study included ECL rabbit IgG, HRP-linked F(ab’)_2_ fragment (Amersham GE, Cat. # NA9340) or ECL Mouse IgG, HRP-linked whole Ab (Amersham GE, Cat. # NXA931) and they were diluted in 5% milk from 1:5000 to 1:10,000.

### 2.9 Pulldown Assay to Test Binding Activity

MBP-His_10_-FAM210A-dMTS was first bound to equilibrated amylose resin (NEB, Cat. # E8201) for 1 hour at 4°C. In addition, a bead-binding control containing amylose resin without adding the fusion protein was prepared. A further control was added where purified MBP was allowed to bind to the amylose resin before adding the cell lysate. Human HEK 293T cells (from one 15-cm culture dish at 90% confluency) were collected and lysed using 500 μl of NP-40 buffer containing 50 mM Tris-Cl pH 7.5, 150 mM NaCl, 1% NP-40, and Roche protease inhibitor. The cell lysate was treated with RNase T1 and DNase I to eliminate the possibility that nucleic acid interactions may affect the FAM210A-EF-Tu interaction. 100 μl of cell lysates were added to the control tubes and FAM210A-containing sample. This mixture was then gently mixed overnight at 4°C to allow binding. The binding reactions were carried out in the same volume with the same amount of cell lysates added. Equal amounts (1 μg) of MBP-His_10_-FAM210A-dMTS and MBP were added. Equal amounts (500 μg) of cell lysates were added to the experimental and control reactions. Overall reaction volumes were equalized with the addition of extra wash buffer. These samples were then purified using gravity flow purification. The beads were washed with wash buffer containing 20 mM Tris-Cl pH 7.5, 200 mM NaCl, 1 mM EDTA, 1 mM DTT, and 1% DDM. Protein samples of the flowthrough, wash, and elution were collected for analysis via Western blot. The volumes of the samples collected were the same (20 μl) for both controls and the experimental sample.

### 2.10 A Negative stain Transmission Electron Microscopy of hFAM210A-dMTS protein

Purified hFAM210A-dMTS was visualized by negative stain transmission electron microscopy. Briefly, three microliters of purified hFAM210A-dMTS (0.005, 0.05, and 0.5 mg/mL) were incubated for 60 seconds on the carbon face of 200 mesh copper grids (Electron Microscopy Sciences) following their glow discharge at 25 mA for 30 seconds. The protein solution was wicked, and the grids were washed 3 times with dH_2_O prior to brief negative staining with a 0.75% uranyl formate. Grids were allowed to air dry and were then imaged using an FEI Talos 120C transmission electron microscope equipped with a CETA-16 CMOS camera with FEI TIA software.

## 3 Results and Discussion

### 3.1 Expression of human FAM210A in different *E. coli* strains

The MBP tag has been shown to increase protein stability and avoid aggregation of select proteins, particularly for transmembrane proteins [9]. Given the property of FAM210A as a transmembrane protein, we selected the His-MBP tag to purify human FAM210A (hFAM210A) protein. The full-length (FL) and mitochondrial targeting signal deletion (dMTS) forms of hFAM210A were cloned into N-terminal His_10_-MBP tagged, or C-terminal MBP-His_6_ tagged expression vectors, 2CT-10 and 2CcT, respectively. Then, the plasmid was transformed into three different *E. coli* strains, BL21(DE3), Rosetta 2(DE3), and Lemo21(DE3), and induced by IPTG for expression trials. In Rosetta (DE3) cells, we did not observe significant expression after IPTG induction for both FL or dMTS FAM210A fused with either N- or C-terminal MBP affinity tag (**Fig. 1A, B**). In contrast, in the BL21(DE3) strain, the dMTS form of FAM210A was strongly induced when fused with the N-terminal His_10_-MBP tag (**Fig. 1A, B**), while the FL form of FAM210A was slightly induced with N-terminal His_10_-MBP tag. In Lemo21(DE3) cells, there was no significant overexpression after induction in the presence of varying L-Rhamnose concentrations (data not shown). The pET28a plasmid (with only His tag) was tested in different strains but failed to overexpress the FAM210A protein (data not shown).

**Fig. 1.**
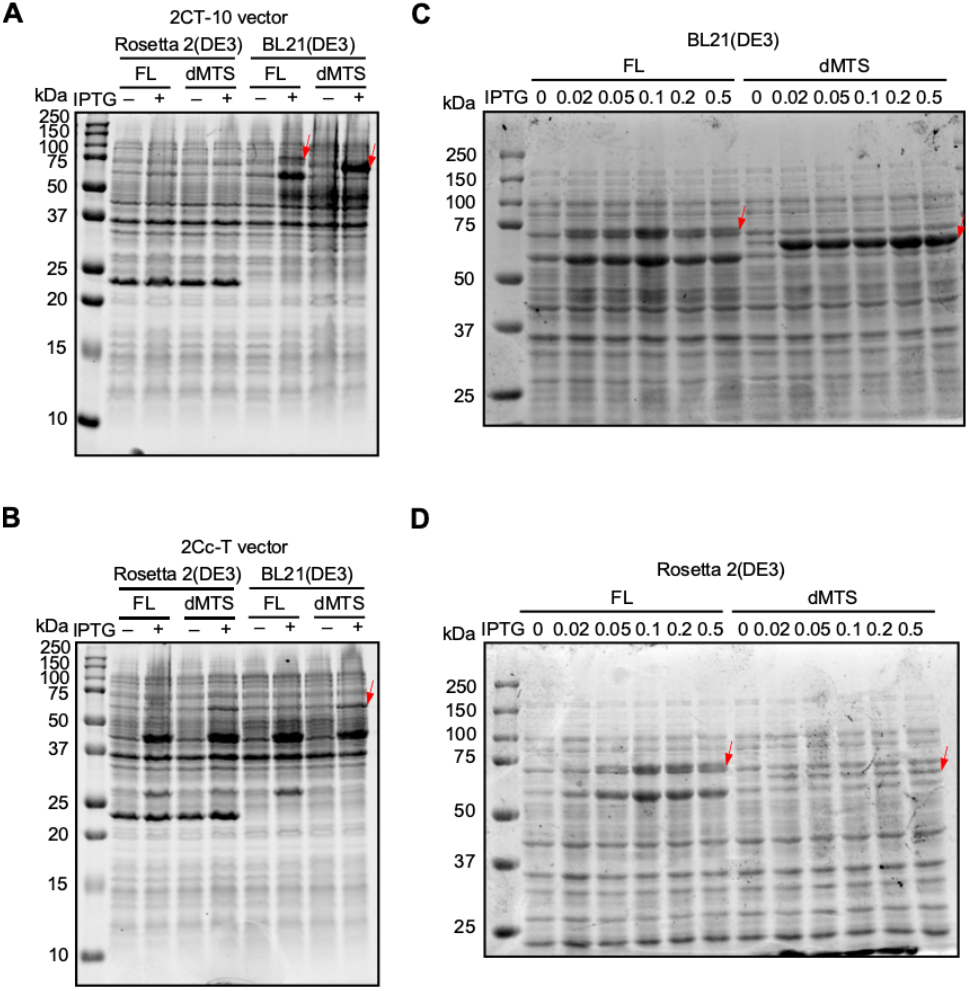
Overexpression of full length and dMTS forms of FAM210A using multiple plasmids, cell types, and IPTG concentrations. (A) 2CT-10 plasmid overexpression trial. (B) 2Cc-T plasmid overexpression trial. (C) IPTG titration trial in BL21(DE3) strain. (D) IPTG titration trial in Rosetta 2(DE3) strain.

Overexpression was also attempted at 16°C for 16 hours to increase the expression levels further compared to previous trials conducted for a shorter time at 37°C. IPTG concentrations ranging from 0.02 mM to 0.5 mM were all tested at this temperature, with no noticeable differences between the samples (data not shown). The FAM210a-dMTS MBP fusion protein was expressed at higher amounts than the full-length fusion protein but still did not reach the levels of overexpression found in samples induced at 37°C for 3.5 hours.

### 3.2 Purification of dMTS form of human FAM210A

Initial attempts to purify the protein from *E. coli* from the soluble, cytosolic fraction were unsuccessful. As FAM210A is a membrane protein, we hypothesized that due to the similarities in mitochondria and *E. coli, the* protein might have been inserted into the *E. coli* cytosolic membrane to be more stabilized during overexpression. We observed that the majority of FAM210A-dMTS MBP fusion protein was localized in the membrane fraction isolated from *E. coli* (**Fig. S1**). To further explore this hypothesis, we isolated the membrane fraction through ultracentrifugation. Following isolation, Triton X-100 and DDM were screened as membrane solubilization agents. Both Triton X-100 and DDM could solubilize the protein; however, given that Triton X-100 complicates the A_280_ detection of protein, DDM was chosen as the detergent in all subsequent trials.

Standard nickel affinity chromatography was used to purify the FAM210A-dMTS MBP fusion protein obtained from the membrane fraction. Following the first affinity chromatography step using Ni-NTA resin, many *E. coli* protein contaminants were observed using SDS-PAGE (**Fig. 2, 4**). A low concentration of imidazole was added to the wash buffer to remove some contaminant proteins that may bind weakly to the nickel resin. Even after this, there are still large amounts of contaminant bands. The total protein yield measured using a bicinchoninic acid (BCA) assay was approximately 1 mg of protein per 1 L of *E. coli* culture.

**Fig. 2.**
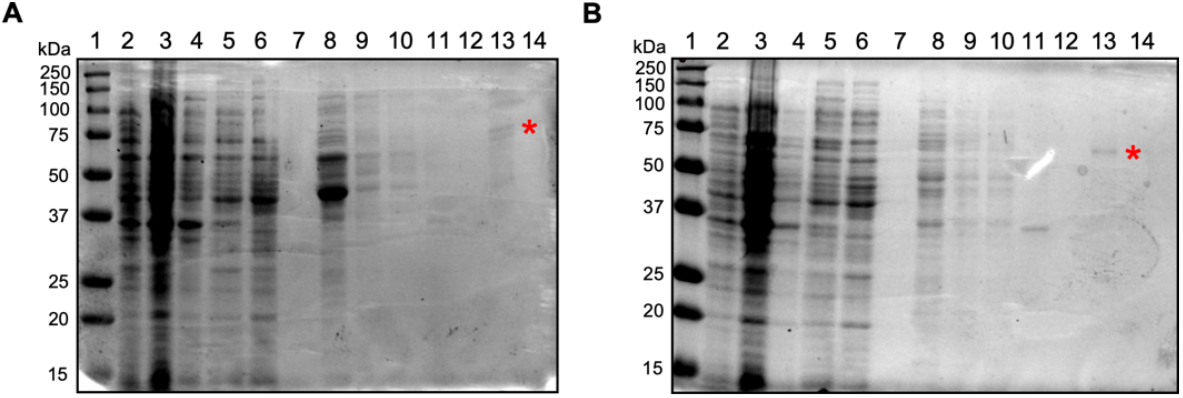
Initial 100 mL purification trials. (A) Purification of full length FAM210A fusion protein from BL21(DE3) cells using a 0.25 mM IPTG induction for 3.5 hr at 37°C. Lane 1: Protein marker ladder. Lane 2: Uninduced cells. Lane 3: Induced cells. Lane 4: Cell debris pellet. Lane 5: Cell lysate flowthrough. Lane 6: Cell lysate supernatant. Lane 7: Cell lysate wash. Lane 8: Elution 1. Lane 9: Elution 2. Lane 10: Elution 3. Lane 11: Solubilized membrane flowthrough. Lane 12: Solubilized membrane wash. Lane 13: Membrane elution 1. Lane 14: Membrane elution 2. (B) Purification of dMTS FAM210A fusion protein using BL21(DE3) cells (100 mL) with 0.25 mM IPTG induction for 3.5 hr at 37°C. * is placed at the solubilized FAM210A fusion protein that was extracted from bacterial membranes. Lane 1: Protein marker ladder. Lane 2: Uninduced cells. Lane 3: Induced cells. Lane 4: Cell debris pellet. Lane 5: Cell lysate flowthrough. Lane 6: Cell lysate supernatant. Lane 7: Cell lysate wash. Lane 8: Elution 1. Lane 9: Elution 2. Lane 10: Elution 3. Lane 11: Solubilized membrane flowthrough. Lane 12: Solubilized membrane wash. Lane 13: Membrane Elution 1. Lane 14: Membrane elution 2.

The second step of affinity chromatography was required to remove the MBP-His_10_ tag from the fusion protein following the cleavage using TEV protease (**Fig. 2, 4**). The nickel resin was used to bind and target the tag and the protease while allowing the FAM210A to be collected in the flowthrough during the purification process. Moreover, the second step of affinity chromatography removed more contaminant proteins and most of the MBP and TEV protease from the sample. To increase the yield of the FAM210A during this step, after the flowthrough was collected, the resin was washed with an appropriate buffer containing no imidazole to collect any protein that may not have been included in the initial flowthrough. This larger volume of flowthrough could then be concentrated using a centrifugal filter. The main contaminant identified in the protein sample can be found at ∼40 kDa in size. This was initially believed to be primarily excess MBP not removed during the affinity chromatography step.

To further increase the purity of the protein sample, size exclusion chromatography (SEC) was attempted to remove the majority of the contaminant proteins. This ultimately proved unsuccessful, as no clear peak correlating to FAM210A existed. Instead, upon analysis of collected fractions, FAM210A was found in all samples, along with other contaminant bands that were not separated (**Fig. S2**). Further analysis of the amino acid sequence for FAM210a-dMTS gave us a theoretical pI of ∼9.1. As this protein is positively charged at neutral pH, ion exchange chromatography was used as an additional purification step (**Fig. 3, 4**). This step was successful in removing almost all of the contaminating proteins; however, the main contaminant band at ∼40 kDa remained. MBP has the opposite charge of FAM210A, so it should have been removed during the cation exchange step, possibly suggesting that the contaminant band was another protein. When starting with 1 L of *E.coli* cell culture, the fusion protein yield is ∼1 mg (Fig. 4, lane 1). Following TEV cleavage, the yield of the cleaved FAM210A-dMTS is ∼0.3 mg (Fig. 4, lane 2). Following ion exchange chromatography, the final yield of FAM210A-dMTS is ∼0.2 mg (Fig. 4, lane 3). The final purity of FAM210A-dMTS protein is around 80% based on Image J quantification.

**Fig. 3.**
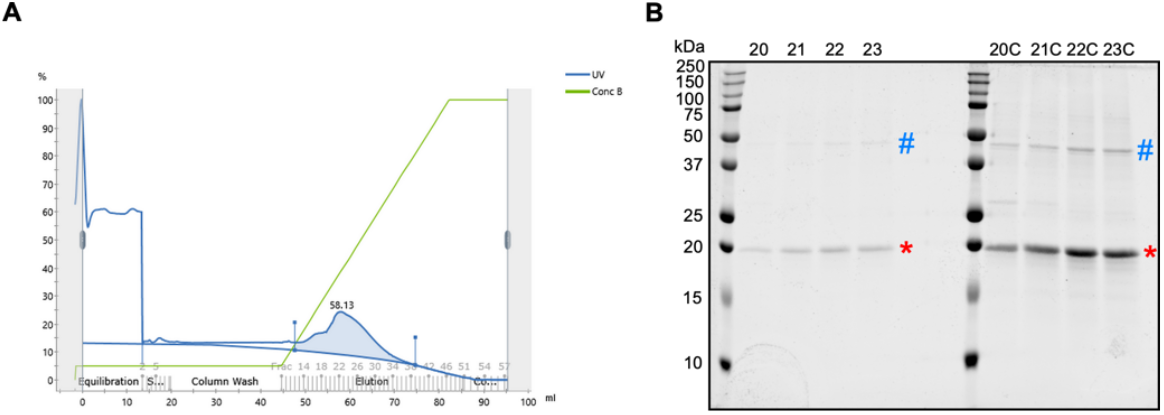
Ion Exchange Chromatography for FAM210A-dMTS purification. (A) A representative chromatogram produced during ion exchange chromatography. A280 absorbance is shown as a blue line. The salt concentration gradient is shown as a green line. The FAM210A containing peak is highlighted in blue. (B) An SDS-PAGE of the unconcentrated and concentrated FAM210A-containing peak samples. C: concentrated. * is placed at the purified FAM210A-dMTS protein. # is placed at a contaminant FAM210A-associated protein band.

**Fig. 4.**
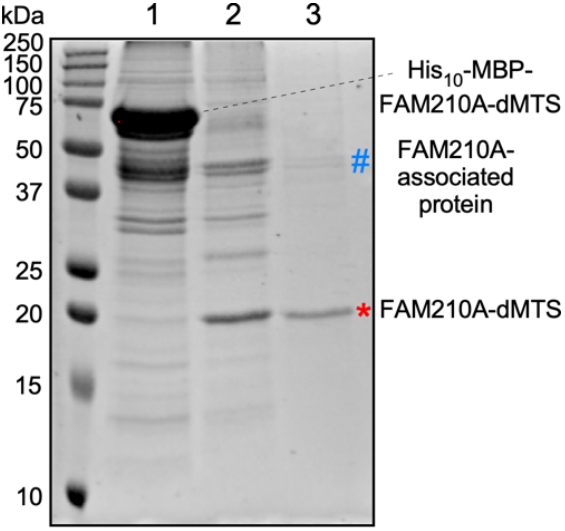
SDS-PAGE gel showing the results of each purification step. Gel was stained with Coomassie G-250 to visualize proteins. Both the fusion protein, the cleaved FAM210A-dMTS (*), and a contaminant FAM210A-associated protein (#) are labelled on the image. Lane 1: first-step nickel affinity chromatography. Lane 2: TEV cleavage and second-step nickel affinity chromatography. Lane 3: Ion exchange chromatography.

### 3.3 Pulldown Assay to Test Protein Binding Activity of FAM210A

A crude pulldown assay was performed to determine whether purified FAM210A-dMTS was active and could still actively bind to a known binding partner. The protein EF-Tu, an essential mitochondrial translation elongation factor, was found to be a top binding partner of FAM210A based on immunoprecipitation (IP) followed by mass spectrometry (MS) and western blot analysis of isolated mitochondria from mouse hearts [8]. Due to this, EF-Tu was chosen as the target for the pulldown assay (**Fig. 5**).

**Fig. 5.**
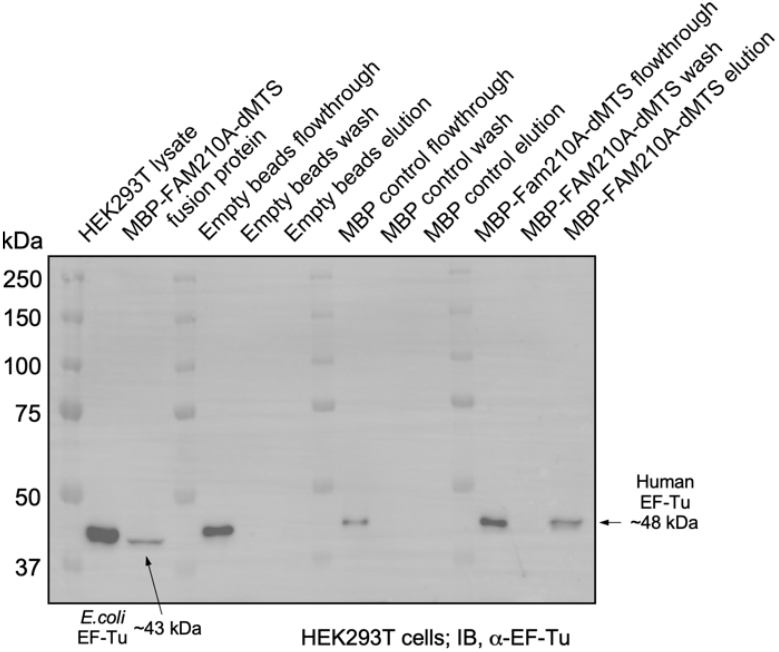
Western blot analysis of the pulldown assay to test binding ability of the purified FAM210A fusion protein. Assay was performed with cellular lysates of HEK 293T cells. The band at ∼48 kDa corresponds to mammalian EF-Tu while the band at ∼43 kDa corresponds to bacterial EF-Tu.

Western blot analysis showed that the fusion protein could bind to the human EF-Tu found in the HEK293T lysate based on a distinct band at ∼48 kDa corresponding to human EF-Tu. The EF-Tu antibody is raised against amino acids 171-455 mapping at the C-terminus of EF-Tu. Due to the high conservation in the antibody-targeting antigen region between humans and *E. coli*, it could bind to bacterial EF-Tu, which can be seen with a band at ∼43 kDa in the fusion protein sample (**Fig. 5**).

The empty resin control elution showed no presence of either mammalian or bacterial EF-Tu, suggesting that the amylose resin could not bind to EF-Tu nonspecifically. Therefore, any EF-Tu present in the experimental sample was pulled down by the FAM210A fusion protein bound to the amylose resin. Furthermore, the additional control using MBP-bound resin showed no EF-Tu binding, eliminating the possibility that EF-Tu interacted with the fusion protein’s MBP tag in the experimental pulldown sample. This experiment was carried out using mouse C2C12 cells, and the MBP fusion protein could also pull down mouse EF-Tu (data not shown). Moreover, human EF-Tu may outcompete or replace *E.coli* EF-Tu from the MBP-FAM210A-dMTS fusion protein as the binding affinity between human FAM210A and human EF-Tu may be higher than that between human FAM210A and *E.coli* EF-Tu (**Fig. 5**, lane 15).

EF-Tu is a highly conserved protein found in both prokaryotes and eukaryotes and has a role as an elongation factor during translation in bacterial cells or eukaryotic mitochondria. The fact that FAM210A has been shown to bind to EF-Tu *in vivo* [8] and now *in vitro* further suggests that FAM210A may serve a role in regulating mitochondrial translation.

After discovering that bacterial EF-Tu had the same approximate size as the MBP tag, it was decided to test post-ion exchange cleaved samples to determine if the contaminant band was EF-Tu. To test this, samples collected after ion exchange chromatography were run at various concentrations before being transferred and probed for EF-Tu since hFAM210A-dMTS had already been shown to bind to bacterial EF-Tu. The results of this Western blot showed that the contaminant band that appeared at∼43 kDa was likely EF-Tu, while MBP and additional proteins cannot be excluded (**Fig. 6**).

**Fig. 6.**
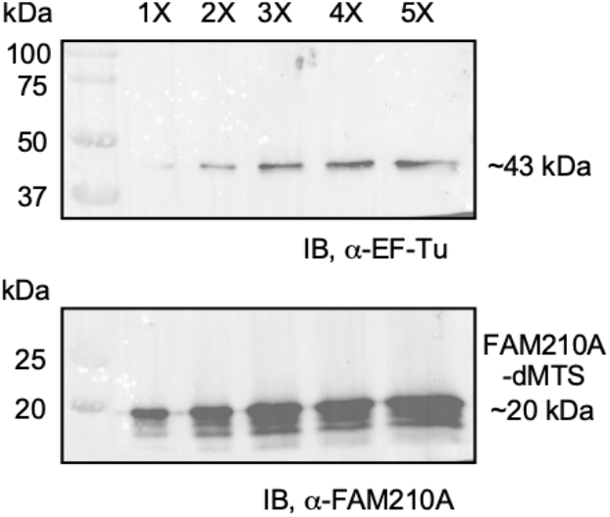
Western blot of samples following ion exchange chromatography. Five different concentrations of the purified sample were run on an SDS-PAGE gel before being transferred and probed for both FAM210A and EF-Tu. A band at ∼43 kDa represents bacterial EF-Tu suggesting that EF-Tu is copurified with FAM210A-dMTS as it is not removed during ion exchange chromatography despite opposing charges.

For future biochemical characterization, removing bacteria EF-Tu (**Fig. 4**, lane 3) would be helpful as it could impair further functional assay. Washing the IMAC and MBP columns thoroughly using a higher concentration of NaCl in the wash buffer for these affinity purification steps might improve the FAM210A purity. Alternatively, identifying the purest fraction of the ion exchange purification followed by a final step on size exclusion chromatography can be further tested for potential structural analysis of the protein.

## 4. Conclusion

In this study, we successfully purified human FAM210A-dMTS from *E. coli*. The purified FAM210A-dMTS fusion protein was shown to interact with known binding partner EF-Tu derived from multiple organisms, including *E. coli*, mice, and humans. Furthermore, the cleaved FAM210A-dMTS could not be separated from bacterial EF-Tu during ion exchange chromatography, showing that the cleaved protein could still interact with EF-Tu (**Fig. 5-6**). *TUFM* (encoding EF-Tu) is evolutionarily conserved in prokaryotes and eukaryotes [11], while *FAM210A* is more recently evolved in *C.elegans* [12] and higher eukaryotic organisms, suggesting that FAM210A is a newly acquired regulatory factor of EF-Tu and mitochondrial translation in multicellular animal organisms.

As much is still unknown about the specific function of FAM210A, this study will allow the simple purification of functional FAM210A-dMTS fusion protein and FAM210A-dMTS for future functional studies and how it may relate to various diseases. Also, the fact that *E. coli* was able to be used as an overexpression and purification system will allow for mutations and changes to be made rapidly to allow for future studies into mutated FAM210A that could be associated with human diseases. This study provides valuable insights into the purification of other mitochondrial membrane proteins in *E. coli* relative to more complex and expensive systems such as insect or mammalian cells.

## ACKNOWLEDGMENTS

We are grateful to Emily Bonanno for her critical reading and polishing of the manuscript. We appreciate the technical assistance from Jermaine Jenkins in ion exchange and gel filtration chromatography and Chad Galloway in CryoTEM analysis. None of the authors have any financial conflict of interest related to the research described in this manuscript.

## Supplemental Figures

**Fig. S1.**
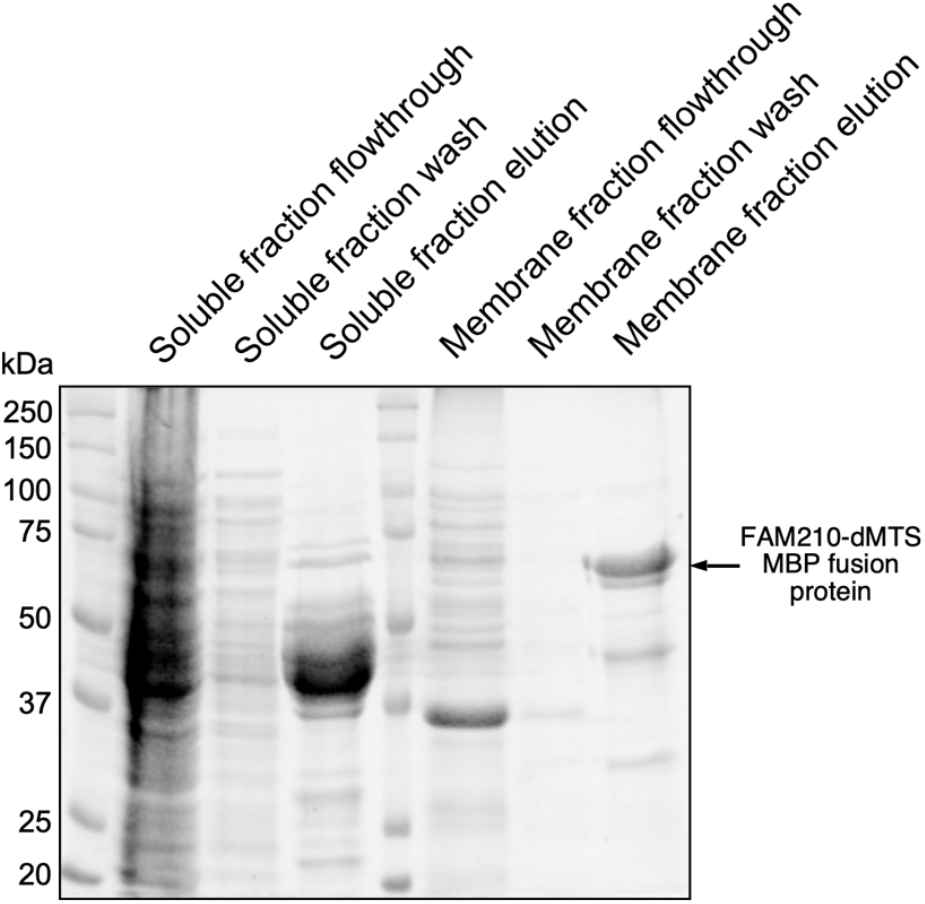
Localization of the FAM210-dMTS MBP fusion protein in the plasma membrane fractions but not in the cytosolic fractions from *E. coli* with overexpression of the fusion protein.

**Fig. S2.**
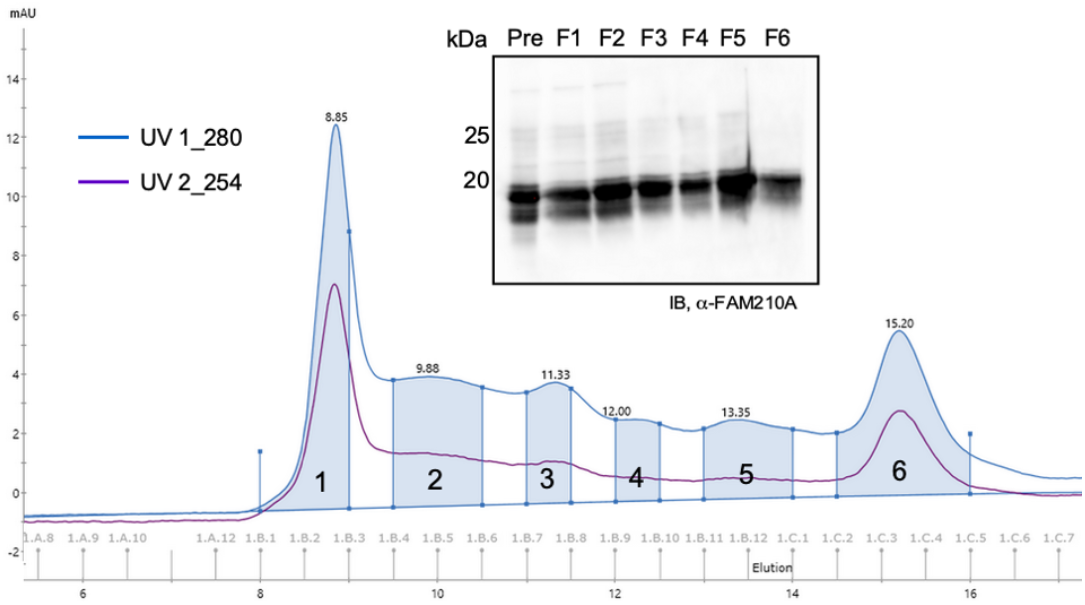
Size exclusion chromatography purification of FAM210A-dMTS followed by Western blot analysis of fractions 1-6.

